# Katydids Shift to Higher-Stability Gaits When Climbing Inclined Substrates

**DOI:** 10.1101/2025.04.04.647231

**Authors:** Calvin A. Riiska, Jacob S. Harrison, Rebecca D. Thompson, Jaime Quispe Nina, Geoffrey R. Gallice, Jennifer M. Rieser, Saad Bhamla

## Abstract

When terrestrial organisms locomote in natural settings, they must navigate complex surfaces that vary in incline angles and substrate roughness. Variable surface structures are common in arboreal environments and can be challenging to traverse. This study examines the walking gait of katydids (Tettigoniidae) as they traverse a custom-built platform with varying incline angles (30°, 45°, 60°, 75°, 90°) and substrate roughness (40, 120, and 320 grit sandpaper). Our results show that katydids walk more slowly as the incline angle increases and as katydid mass increases, with a decrease of around 0.3 BL/s for every 1° increase in incline. At steeper inclines and larger sizes, katydids are also less likely to use an alternating tripod gait, opting instead to maintain more limbs in contact with the substrate during walking. Katydids also increased average duty factor when climbing steeper inclines and with increasing body mass. However, substrate roughness did not affect walking speed or gait preference in our trials. These findings provide insights into how environmental factors influence locomotor strategies in katydids and enhance our understanding of effective locomotor strategies in hexapods.

## 1 Introduction

Rainforests are ecosystems that feature substrates with varying inclines and surface roughnesses. For many species inhabiting these regions, mastering effective movement techniques on highly variable surfaces while combating gravity is vital for survival; otherwise, they face the severe risks associated with falling from heights. Many organisms, including vertebrates and invertebrates, have evolved traits to assist in arboreal locomotion. Some organisms use specialized body structures, such as elongated limbs [1, 2], adhesive pads [3, 4, 5, 6, 7], or prehensile tails [8]. While others shift their underlying behavior, such as balancing maneuvers [9, 10, 11], redistribution of leg drive [12], changing limbs that support weight [13], or cyclic movements of the center of mass to increase stability [14, 15]. Examining how and when organisms change their movement behavior during climbing will shed light on their interactions with the environment and their ability to navigate complex and challenging environments.

Insects are commonly observed using an alternating tripod gait while walking over a substrate. This movement pattern involves three limbs contacting the ground at any one moment: the front and hind limbs on one side, and the middle leg on the opposite side (Fig. 1a). This limb configuration creates a support polygon which typically contains the center of mass [16]. This gait is symmetrical, with each trio of limbs being highly coupled and, as has been shown in *Blaberus* cockroaches, functioning like a leg of a biped [17, 18] as they are able to produce body dynamics common to runners with various numbers of limbs despite each limb uniquely and independently influencing the acceleration [19, 20]. The tripod gait is considered highly energy efficient [21] and is often regarded as one of the fastest methods for hexapod movement [22] with energy and ground reaction forces resembling running or bounding gaits of other animals at high speeds [18, 17]; other arthropods have been observed using intriguing galloping gaits [23] and metachronal gaits with reduced vertical oscillations [24] at high running speeds.

**Figure 1.**
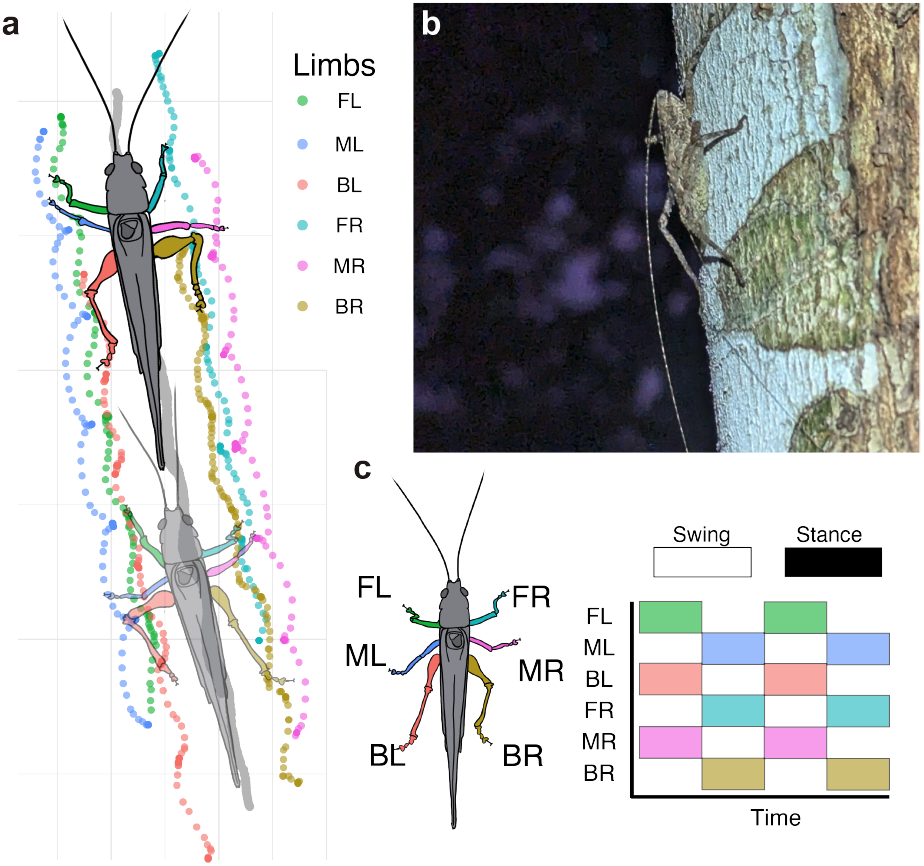
(A) A diagram of a katydid using an alternating tripod gait to traverse a substrate. (B) A katydid in the Peruvian Amazon Rainforest maneuvering across a tree trunk. (C) This panel represents an idealized alternating tripod gait when hexapods, such as katydids, move in an alternating tripod gait; the front left, middle right, and back left limbs coordinate their movements. Colors represent individual limbs in the gait plot, indicating when they make contact with the substrate. The swing phase denotes when a limb is in motion, while the stance phase represents when the limb maintains contact with the substrate.

We aim to investigate gait transitions in arboreal katydids (family: Tettigoniidae) found in the Peruvian Amazon. Little is known about the specific gait mechanisms of katydids. Many species in this family on multiple continents are highly arboreal [25, 26, 27, 28], navigating through diverse and complex terrain. Though limb patterns may deviate from a classic tripod gait due to longer-than-normal hind limbs [29], a closely related species within the orthopteran family has been shown to use the tripod gait [30], which may be advantageous for the challenging terrains that katydids navigate. A major case where the tripod gait has been successful on rough, smooth, and inclined terrain is when it is employed by cockroaches. By slightly altering their strides [31] through a mechanical feedback system that stabilizes the body when they run rapidly [32], they are easily able to overcome uneven terrain. The stability of the tripod gait allows cockroaches to traverse a wide variety of challenging substrates [33] and, combined with adhesive footpads, perform acrobatic feats overcoming rapid changes in incline [9]. Behavioral experiments [16] and computational models [22] have shown that the tripod gait remains stable on both horizontal and vertical surfaces [34], and indeed, the tripod gait has been observed in climbing cockroaches [14, 9]. When cockroaches climb, their front legs pull the head toward the wall while the hind legs push the abdomen away, and they maintain this posture. All legs also pull inward toward the midline, a force pattern shared by geckos. These results suggest that limbed animals use a common template for climbing and hexapods may fare well using a tripod gait [14].

On uneven terrain, factors such as incline angle and surface roughness can significantly impact an organism’s speed and gait. For instance, in ants, rough surfaces can influence running speed by affecting adhesive and propulsive forces [35, 36] with smaller species slowed more than larger species as surface features increase in size [37]. An increase in surface incline generally reduces running speed, alters the direction of the forces exerted by the limbs due to changes in the ground reaction force, and may modify the insect’s gait [35, 38, 13, 14]. In natural settings, incline and roughness often vary across different spatial and temporal scales, forcing organisms to navigate diverse, challenging substrates. Understanding how and when organisms adjust their kinematics and behavior in response to changes in substrate can offer insights into how animals navigate and thrive when encountering complex features in their environments.

This study investigates how surface incline and roughness affect the climbing behavior of katydids. We conducted field research at the Finca Las Piedras research station in the Peruvian Amazon over a period of 10 days as part of the in-situ Jungle Biomechanics Lab stupski2024curiositydrivensciencesitujungle. During this time, we observed several katydid species climbing trees and positioned on inclined surfaces that varied from smooth to rough textures (Fig. 1b). This led us to focus on this clade further. We hypothesize that steeper inclines and smoother surfaces will complicate climbing in katydids, resulting in reduced climbing speed, though they will maintain a tripod gait. This research aims to fill knowledge gaps related to the biomechanical and ecological aspects of gait transitions in insects when faced with substrate changes that may occur within their natural environments.

## 2 Methods

### 2.1 Specimen Collection and Experimental Protocol

Over a period of 10 days in August 2024, we collected 24 katydids from the Peruvian Amazon Rainforest at the Finca Las Piedras Research Station (Parcela 4, Sector Monterrey, Peru; - 12.225718266152889, -69.11430715640181). Individuals were collected at night using a hand net and then transferred into foldable mesh insect containers. All katydids were imaged, underwent experimental trials and were kept in captivity for up to two days; they were then released at their collection site. Each individual was marked with white, non-toxic paint on their head to assist in video tracking and to ensure no recapture of individuals (Collection Permit: 1288-2022-GOREMAD-GRFYF).

We constructed a custom-designed walking platform measuring 22 cm by 28 cm from aluminum extrusion parts (Fig. 2a). The walking platform was adjustable and could be set to five different inclines: 30, 45, 60, 75, and 90 degrees. We could also adhere various slips of sandpaper to the surface to adjust the roughness that the katydid moved across. We used three different grit types of sandpaper from McMaster-Carr: 40, 120, and 320 (22 cm by 28 cm). A GoPro Hero 10 Black camera was mounted 36 cm away, positioned orthogonally to the walking surface to record the walking trials and filming at 60 Hz.

**Figure 2.**
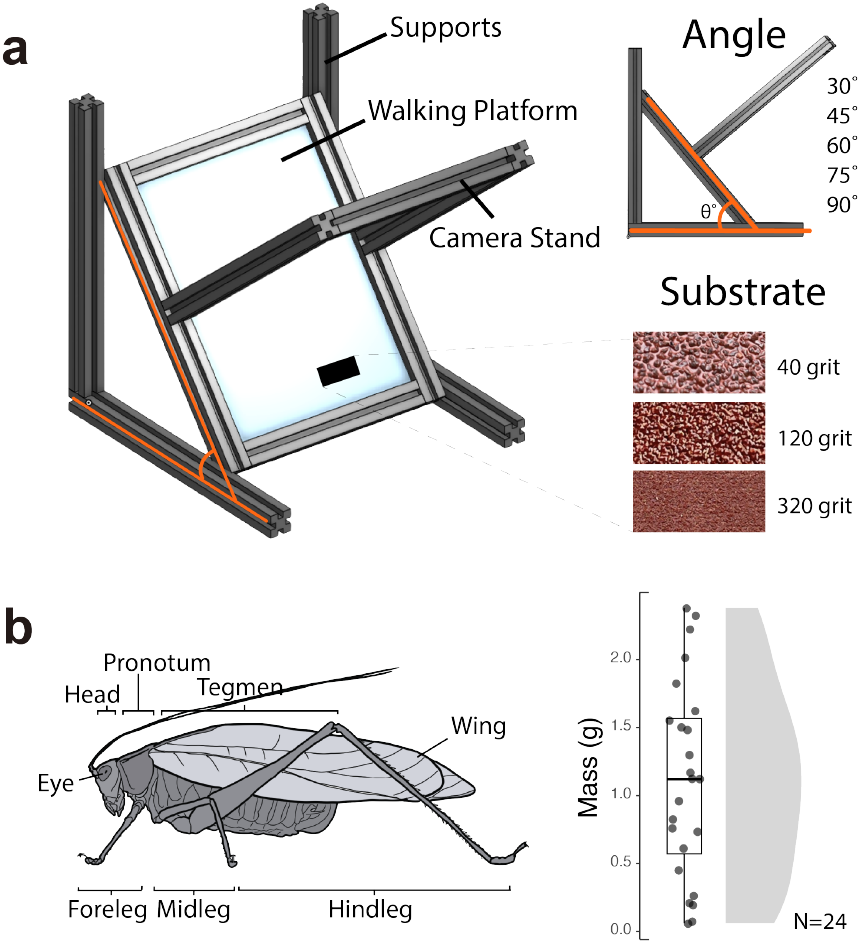
(A) A schematic of the experimental climbing wall. The wall was designed so that the incline and frictional substrate could be modified for different experimental trials (S.Video 1). (B) A drawing of the lateral view of a typical katydid with its limbs and body sections labeled. Katydids collected for this dataset weighed between 0.06 and 2.38 grams. Points represent individual katydids, while the shaded region illustrates the distribution.

During each trial, an individual was positioned at the bottom of the inclined platform and allowed to climb upward. If no movement occurred, the individual was gently nudged from behind. If the katydid still did not ascend, the trial was restarted. Ideally, each individual would complete five sequential runs for every combination of incline and substrate, presented in random order. However, due to fatigue or lack of incentive, not all individuals were able to complete all trials. Trials in which the katydid failed to secure a grip or did not move were excluded. In total, the 24 individuals collectively completed 492 trials, with each individual contributing between 2 and 39 trials (Mean: 20.5 *±* 11.4 trials). Each individual experienced between 1 and 9 distinct combinations of incline and surface roughness. A complete list of trial numbers, along with the grit and angle combinations for each individual, is provided in the supplement (S.Fig. 1 and 2), accompanied by a power analysis to quantify the observed power of the experiment(S. Fig. 3).

### 2.2 Tracking and Gait Analysis

To measure body speed, the white paint marked on each katydid was automatically tracked using a custom-written MATLAB code. This process involved isolating the katydid by subtracting median image pixels, binarizing the image, dilating and eroding to isolate the white dot. We then tracked its position over time to generate body position data and used the `peakVals` function in MATLAB to find the local maxima of body velocity. Pixels from videos were calibrated to the millimeter scale using a ruler. We then averaged across the local maxima to obtain an average of the peak velocity during a walking trial for each individual. We use the local maxima of speed instead of a full average of speed because we did not want pauses in the motion between strides to influence the data. We then averaged trials within individuals for repeated measures on the same grit and angle combination and used those averages for statistical analysis (described below). All body speed was converted into body length per second (BL/s) using the combined length of the head, pronotum, and tegmen (Fig. 2b).

To look at gait transitions across grit and angle, we behaviorally scored whether an individual used an alternating tripod gait (the front and hind legs on one side and the middle leg on the opposite side move simultaneously; S. Video 1) or used a different gait (with 4 or more limbs in contact with the surface at one time; S. Video 2) during their upward climb for each trial. Each trial was reviewed twice, once each by two separate observers (CAR and JSH). The trial was then marked with either a tripod or non-tripod gait. We averaged trials within individuals for repeated measures on the same grit and angle combination to determine a percent use of tripod gait for that individual on that particular incline and grit combination. Additionally, we compared observational scoring to duty factor (explained below) to compare observations against a quantitative metric for gait biomechanics (S. Fig. 4). We used those averages for statistical analysis (described below).

A random subset of videos (45 in total from 19 individuals) was manually tracked using `DLTDV8` in MATLAB [39] for a more detailed analysis of limb coordination, stride length, and stance duration. For each trial, we tracked frame-by-frame the anterior-most tip of the head, the posterior tip of the abdomen, and each limb during a trial and then calculated the displacement of the points across time normalized by body length using a custom-written code in R (RStudio, v2024.09.1). We calculated the first derivative with respect to time and then smoothed the resulting speed traces with a third-order Savitzky–Golay filter over an 11-frame window. We defined a limb to be in swing if its smoothed speed was above 0.015 body-lengths per frame; otherwise, it was labeled as in stance. We calculated the duty factor of each limb to be the proportion of frames spent in stance. For each swing event, we then summed the displacement of the limb relative to body length to yield a stride length. We then averaged across all limbs to get an average duty factor, stance duration, and stride length for each individual for each trial (S.Fig. 5).

### 2.3 Statistical Analysis

To look at how the incline angle and substrate affected walking speed, we built a linear mixed-effects model (LMM) in R using the `lmer`. We converted the angle into a centered, numeric predictor and included individual mass as a covariate. As fixed effects, we included the incline angle, the sandpaper grit, and body mass, and we tested both all two-way interactions and then a full three-way interaction. A parameter that represented the individual was included in the model as a random effect to account for repeated measures and inter-individual variation. We compared the two-way versus three-way formulations using a likelihood-ratio test, and then tested whether adding a random slope for angle improved the fit, which it did not. P-values for each fixed effect were obtained via Satterthwaite’s method. Finally, we assessed whether a random-intercept-only structure sufficed and used the `MuMIn::dredge()` function on the full three-way model to confirm that the two-way model (with no random slope) was best supported by AICc (Table 1; S.Table 1-3; S. Fig. 6).

**Table 1.** Fixed-effects estimates from the Linear Mixed Model predicting peak speed (BL/s).

| Term | Estimate | Std. Error | df | t-value | p-value |
| --- | --- | --- | --- | --- | --- |
| (Intercept) | 2.17 | 0.25 | 22.39 | 8.57 | $1.62 \times 10^{-8} *$ |
| Grit | -5.01 | 11.17 | 80.93 | -0.45 | 0.67 |
| Angle | -0.30 | 0.07 | 81.80 | -4.19 | $6.93 \times 10^{-5} *$ |
| Mass | -2.08 | 0.36 | 22.46 | -5.82 | $6.93 \times 10^{-6} *$ |
| Grit:Angle | -13.11 | 8.22 | 79.73 | -1.60 | 0.11 |
| Grit:Mass | -18.75 | 16.98 | 80.49 | -1.10 | 0.27 |
| Angle:Mass | 0.20 | 0.10 | 82.36 | 1.98 | 0.05 |

To determine whether the incline angle or substrate affected the usage of the tripod gait, we built a generalized linear mixed model (GLMM) using the `glmer` function in R. The response variable was the proportion of tripod gait usage, modeled as a binomial outcome. Similar to the body speed LMM, we used the incline angle, sandpaper grit, and body mass as fixed effects, and we first compared a two-way model to a three-way model via a likelihood-ratio test. Since the three-way interaction was not significant, we retained only two-way interactions. The individual ID was included as a random effect. Next, we tested whether adding a random slope on angle significantly improved fit, which it did. We fitted the full factorial two-way model set and identified the best fit using their AICc scores (Table 2; S.Table 4,5; S. Fig. 7). Because we detected under-dispersion, no observation-level random effect was added. To evaluate our best fit models, we examined the residual versus fitted plots.

**Table 2.** Fixed-effects estimates from the Generalized Linear Mixed Model predicting tripod gait usage.

| Term | Estimate | Std. Error | z-value | p-value |
| --- | --- | --- | --- | --- |
| (Intercept) | 0.82 | 0.27 | 3.02 | 0.0025 * |
| Grit | -7.48 | 21.03 | -0.36 | 0.72 |
| Angle | -1.77 | 0.32 | -5.57 | $2.61 \times 10^{-8}$ * |
| Mass | -1.35 | 0.38 | -3.59 | 0.00033 * |
| Grit:Angle | -4.52 | 18.61 | -0.24 | 0.81 |
| Grit:Mass | 29.37 | 34.62 | 0.85 | 0.40 |
| Angle:Mass | -0.76 | 0.43 | -1.76 | 0.079 |

## 3 Results

Species identification from the 24 katydids used in this study indicated that we had individuals from approximately seven genera across three Supertribes of Tettigoniidae (Pseudophyllinae, Phaneropterinae, and Conocephalinae). Unfortunately, we were unable to export specimens or collect tissue samples, which prevented us from identifying the species level. Nonetheless, individuals were identified to at least the subfamily level using dorsal images and confirmed with assistance from a taxonomist specializing in Peruvian katydids (S. Note 1). Individuals in the study ranged from 0.06 to 2.38 grams (mean: 1.11 *±* 0.72 g) and from 0.83 to 6.89 cm (mean: 4.3 *±* 1.9 cm; Fig. 2b). Body speeds varied from 0.2 BL/s (Grit:40, Incline Angle: 75) to 12.2 BL/s (Grit:320, Incline Angle:30), translating to absolute distances of 1.3 to 13.3 cm/s (mean: 6.6 *±* 3.0 cm/s; Fig 3). The average stance duration of katydids was between 0.13 and 1.6 seconds (mean:0.48 *±* 0.36 s). The average stride length of katydids across the trials was 0.243 to 1.309 BL (mean: 0.676 *±* 0.13 BL). However, stride length decreased significantly with increasing body mass (F-stat= 29.2, adj. R^2^ = 0.4, p= 2.9 *×* 10^*−*6^; S. Fig. 5). The percentage chance of using an alternating tripod gait among individuals ranged from 42.7 to 100 percent, although the likelihood of using this gait depends on multiple factors (see results below).

During experiments, katydids only fell from the wall in two combinations of grit and incline angle (120 grit: 90° incline and 120 grit:60° incline). During failed trials, katydids fell from the wall in 16% (4 of 24) and 12% of trials (2 of 17) with a grit of 120 and angles of 90° and 60°, respectively. Unfortunately, we were unable to collect any trials from katydids climbing on 320grit:90° incline (Fig. 4C). Trials in which katydids failed to climb were not included in the analysis for body speed or gait. We found no trends relating the failure rate to individual or body size.

**Figure 3.**
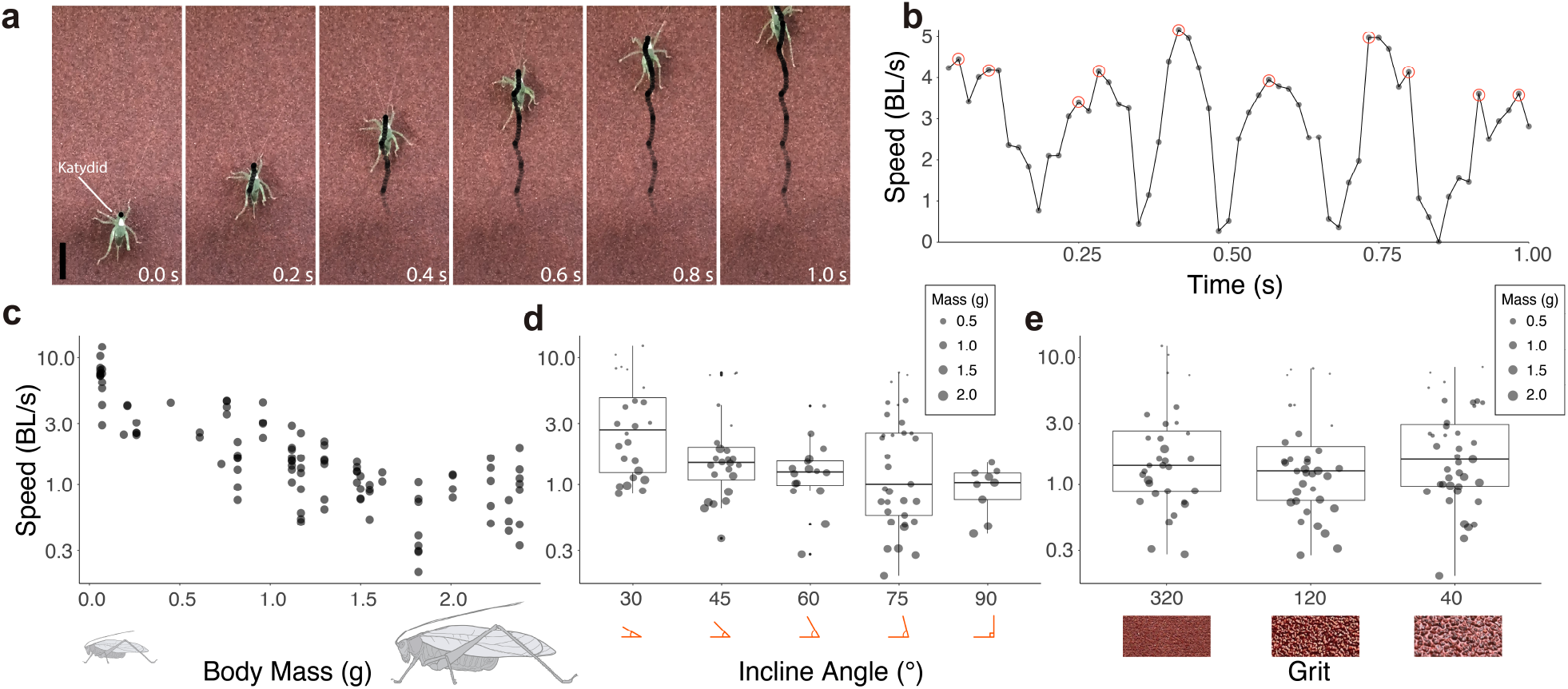
(A) Image sequence from GoPro footage showing a katydid traversing the climbing wall over time. The black points represent the tracked points used to calculate body speed. The vertical black line serves as a scale bar of 2 cm. (B) Body speed for a characteristic trial of a katydid walking experiment. Points indicate the tracked body speed between frames. Red circles highlight instantaneous peak speeds used to calculate the average peak speeds of katydid locomotion. (C) Average peak speed (BL/s) relative to body size. Each point represents the average speed of an individual within a specific incline angle and grit combination. (D) Average peak speed (BL/s) with respect to incline angle. (E) Average peak speed (BL/s) with respect to substrate roughness. For panels D and E, points represent the average speed of an individual within a specific incline angle and grit combination, and point size indicates the mass of the individual.

**Figure 4.**
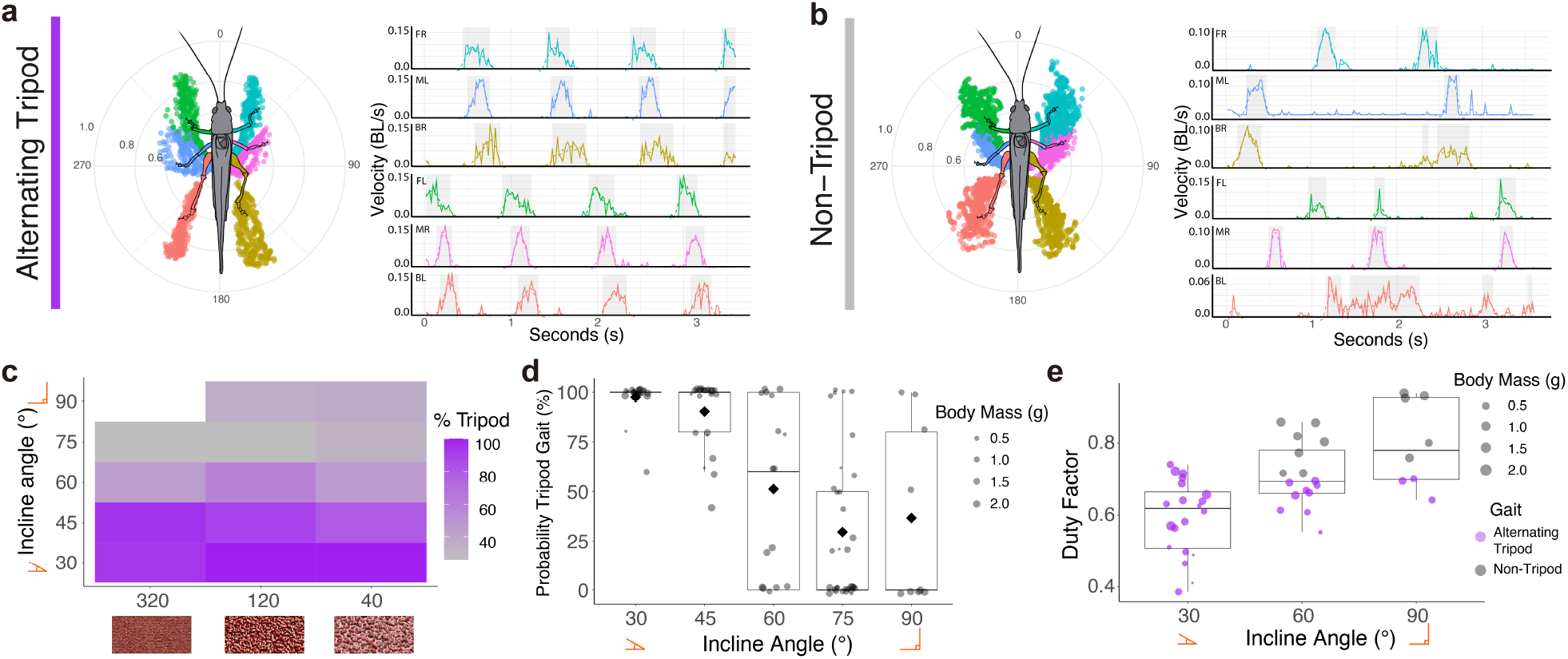
(A) A characteristic alternating tripod gait of a katydid is first shown via a circular plot where points represent limb motion relative to body position during a trial. The velocity plots next to the circular plot indicate the speed in body lengths per second of each limb during the trial. (B) A characteristic non-tripod gait from a katydid is shown via a circular plot where points represent limb motion relative to body position during a trial. The velocity plots next to the circular plot indicate the speed in body lengths per second of each limb during the trial. In panels A and B, the solid line shows raw data from frame-by-frame tracking, and the shading indicates when a limb was classified as in a swing phase. The colors of the points and lines correspond with specific limbs. In the circular plots, distances and angles are relative to the body’s centroid. (C) A heatmap displays the percentage of tripod gait usage across different inclines and grit levels, with color indicating the percent change across trials. (D) The probability of tripod gait across incline angles is shown, with point size indicating individual mass. The black diamond represents the average percentage of individuals using a tripod gait at each incline angle. (E) The duty factor, calculated from manually tracked trials, increases on steeper inclines. Points indicate the average duty factor for an individual on that particular slope, with point size reflecting the organism’s mass. Colors correspond to behavioral scoring of alternating tripod or non-tripod gait.

The best-performing model in our linear mixed-effects model (LMM) of walking speed included incline angle, body mass, and grit as fixed effects, along with their two-way interactions (Table 1). The three-way interaction did not improve the fit, and adding a random slope for angle was not justified; thus, the final model retains only the two-way interactions and a random intercept for individuals. In the model, peak speed decreased with increasing incline angle (Estimate = *−* 0.301, *t* = *−*4.19) and decreasing body mass (Estimate = *−*2.075, *t* = *−*5.82), while grit on its own was not significant (Estimate = *−*5.006, *t* = *−*0.45). The only borderline interaction effect was between angle and mass (Estimate = 0.199, *t* = 1.98, *p* = 0.051), suggesting that heavier individuals slow down slightly less at steeper angles. The variance for the random intercept was estimated at 1.29, indicating substantial individual-level differences in baseline walking speed.

In our generalized linear mixed-effects model (GLMM), the best-performing model (based on AIC) incline angle, body mass, and their interaction (Table 2), with grit included but its interactions deemed non-significant. We did not retain the three-way interaction of grit × angle × mass. Adding a random slope on angle substantially improved the fit. In that model, incline angle had a strong negative effect on tripod gait usage (Estimate = *−* 1.771, *z* = *−*5.57), and body mass also had a negative effect (Estimate = *−*1.349, *z* = *−*3.59). Grit had no significant main effect (Estimate = *−*7.481, *z* = *−*0.36), nor did its interactions with angle or mass. The angle × mass interaction was marginal but not significant (Estimate = *−*0.762, *z* = *−*1.76, *p* = 0.079), suggesting heavier individuals may reduce tripod use slightly more on steeper slopes. Random-effects variance was 0.69 for the individual intercept and 0.86 for the angle slope, reflecting individual differences in both baseline gait choice and how slope alters gait.

The mean duty factor for manually-tracked trials was 0.673, indicating that limbs spent, on average, 67.3% of the gait cycle in the stance phase (min = 0.386, max = 0.939). However, the duty factor increased with both mass (F-stat= 45.2, adj. R^2^=0.52, p= 3.6 *×* 10^*−*8^) and substrate angle (F-stat= 27.1, adj. R^2^=0.38, p= 5.4 *×* 10^*−*6^), which indicates that larger katydids and steeper incline trials lead to longer average stance phases and more stable gaits.

## 4 Discussion

This research investigated the walking behavior of katydids (Tettigoniidae) on a platform with varying incline angles and substrate roughness. Our results indicate that katydids walk more slowly when climbing steeper inclines, and larger katydids also walked more slowly. However, substrate roughness had no noticeable effect on their walking speed. Similarly, gait patterns were affected by both incline and mass, but not substrate. When traversing steeper inclines, katydids were more likely to shift away from using the alternating tripod gait toward more stable gaits: ones with increased duty factors [40], more limbs maintaining contact with the substrate [34], and therefore higher adhesive forces keeping the insects from falling [41, 7, 42]. Notably, the individual effect on body speed and gait was significant in our experiment, suggesting that intrinsic traits within an individual play an important role in locomotion. Nevertheless, despite the strong individual effects, our models demonstrate that body size and incline have a significant impact on locomotor strategies, while substrate roughness does not. Below, we discuss these findings and the limitations of this study in more detail.

Some creatures, including geckos and cockroaches, adeptly navigate inclined and vertical substrates and are capable of rapidly climbing with minimal difficulty [3, 14, 40]. However, in some cases, aligning with our results, increasing incline results in significant variation in body speed and/or movement patterns, particularly when additional challenges like granular surfaces are present. For example, *Componotus* ants alter their movement from a tripod gait to a metachronal gait (where one limb moves at a time) when ascending a slippery, granular slope, reminiscent of an ant lion pit [43]. Additionally, formicine ants exhibit slower walking speeds when subjected to higher slopes accompanied postural changes lowering the body closer to the surface [44, 19] and higher duty factors in an effort to gain stability [40]. Many organisms, including horses [45], humans [46], and cockroaches [17], show clear relationships between gait and speed, where adjusting body speed often involves changing gait patterns. From our experiments, it is not entirely clear whether the gait shift causes the katydids’ slower body speed or if it is just a behavioral response as they adapt to more complex substrates. Future research should investigate this relationship further.

Climbing offers significant ecological advantages, whether by locating resources or escaping predators. Mathematical models addressing attachment mechanics in climbing animals suggest that within taxa, attachment performance to a vertical substrate may be size-independent as attachment pads grow isometrically with animal mass but increase in efficiency [47]. However, phylogenetic studies have demonstrated extreme allometry in the footpad area for vertebrates and arthropods, supporting the mathematical models but indicating a potential limit that could complicate adhesion-based climbing for organisms larger than geckos [48]. In that study, it was also found that hemimetabolous insects, such as katydids, possess smaller relative foot pads than their holometabolous counterparts or vertebrates, which may explain why incline angle influences katydid climbing so strongly.

Surface texture in our experiments did not significantly affect the movement speed or gait pattern of katydids. Claws on the ends of the insects’ tarsi enhance frictional interactions and strongly aid in attachment [49, 50, 51], but this indicates that the friction force is similar on all of the surfaces tested, or it is playing a relatively minor role in this system. Katy-dids also likely rely on the adhesive properties of their tarsi, which enable them to cling to various textured surfaces. The tarsi enhance locomotion upon contact with the ground due to specialized microstructures and fluid-based adhesion. Some insects possess hair-like structures (setae) that conform to small surface irregularities [52, 53, 54], creating numerous independent contacts through van der Waals forces [55, 47]. However, other insects, like katydids, feature smooth pads with embedded fibers that deform around surface features, increasing contact area and enhancing adhesive and frictional forces [53, 7, 56, 57]. Dry adhesion alone is not sufficient to support the insect and is further supported by secretions [52, 58, 59, 42, 54]; insects with setae use internal tubes for fluid delivery, while those with smooth pads secrete it through pores. This secretion boosts pad contact area, adds viscosity, and facilitates capillary adhesion [58]. Although the composition of the secretion remains unclear, similar lipid-like substances are found in footprint droplets. While the secretion is vital for adhesion, structural adhesion is also important [42].

Few studies have quantified how much weight an orthopteran’s tarsus can support. In some species, the tarsus divides into four segments (tarsomeres), with different combinations engaging based on surface orientation [42, 41]. Jiao et al. [42] measured the adhesive force normal to a surface of a tarsomere in *Tettigonia viridissima* (Great Green Bush-cricket), which plateaued at 1.1 mN with loads exceeding 0.8 mN. Since a typical *T. viridissima* weighs about 1 g (9.8 mN), each leg must exert approximately 1.6 mN to support the maximum weight of the insect. The adhesive force from a single tarsomere is inadequate, but engaging multiple tarsomeres can increase adhesion [42]. In another taxonomic group, Labonte and Federle [6] demonstrated that stick insects with adhesive pads increase their adhesion by pulling tangentially, generating greater attachment. If more legs contact the surface, the force requirements for each individual leg decrease, reducing the need for tangential forces [6]. Our results suggest that as the incline angle increases, katydids move more slowly and shift away from the tripod gait, presumably to keep more feet in contact with the substrate and increase the support force they can generate. Future studies could measure the grip strength of Tettigoniidae and determine the angles and masses at which a tripod gait is insufficient for supporting a given mass and whether they use other behaviors, like stick insects, to assist in supporting body mass. Future studies may also look at the direction of forces that each tarsus generates as it is likely that the front legs will act to pull the animal while the hind legs act to push upward and into the surface, with the opposite effect occurring in downward climbs [13].

A key limitation of our study is that we could not focus our data collection on a single katydid species or developmental stage. Preferably, our study would use specimens from within a single species. This limitation stems from the logistical challenges of researching in a remote field environment under time constraints, as well as the relative abundance of katydids. We could not export specimens from the field station, making precise species-level identification impossible. Additionally, managing the motivation of an organism across so many trials is difficult. Although the order of grit and angle combinations was randomly selected for each individual, we could not run each katydid on every grit and incline angle combination. Consequently, variations in morphology and behavior across different species and developmental stages introduce additional variability into our model. This is evident in the large variance in the random intercept for our mixed models. However, our data still indicate that there are constraints on katydid climbing and gait. Even after accounting for individual variance in the models, we observe that incline angle and body mass negatively impact movement speed and overall gait. Future research could explore the allometric scaling of Tettigoniidae foot pads, calculate the safety factor of the katydid during adhesion, and investigate climbing behavior within a single species to better understand precisely why this transition in gait occurs and whether it relates to the need to support body mass while climbing.

This study and related follow-up studies could aid in the design and improvement of hexapod and climbing robots. Hexapod robots, like RHex, enable comparative studies of limb compliance, gaits, and locomotion on varied surfaces [60, 61]. A template for climbing in both hexapods and quadrupeds—where mass is accelerated and anchored periodically—has already inspired robot models for climbing [14]. The hexapedal robot RiSE mimics the splayed stance of climbing geckos and squirrels and follows a similar force pattern through its legs [62]. Stickybot is an example of a climbing robot that uses dry adhesive pads to attach during climbing and is capable of scaling smooth, 90°inclines at a speed similar to RiSE [63]. Future research into how katydids and other hexapods shift their gaits at varying inclines could enhance the design or programmed gait in robots.

## 5 Conclusion

Our research indicates that steeper slopes and larger body sizes negatively impact katydid walking speed and that these factors cause katydids to shift away from alternating tripod gaits. Moreover, we found that there can be marginal interactive effects among substrate variables, which can influence the organisms’ responses. Nonetheless, the substantial individual influences highlighted in our models suggest that species-specific morphology is crucial for locomotor performance. This study offers novel insights into Katydid locomotion in the Peruvian Amazon, enhances our understanding of insect biomechanics on substrates modeling natural environments, and may inform the design of bio-inspired robotic systems.

## 6 Supplementary data

Supplementary data and code are available in Dryad (DOI: TBD) and upon author request.

## 7 Competing interests

No competing interest is declared.

## 8 Author contributions statement

C.A.R., J.S.H, and J.M.R. conceived the experiments. C.A.R., J.S.H., J.Q.N., and G.R.G. assisted in specimen collection and field research. C.A.R. conducted the experiments, and C.A.R., J.S.H., and R.D.T. analyzed the results. J.M.R. and S.B. supervised this research. S.B. secured funding. All authors contributed to writing, discussion, and revising the manuscript.

## Acknowledgments

The authors thank Johana Reyes-Quinteros and the 2024 Jungle Biomechanics Class for their assistance in the field. We also thank Dr. Dave Nickle for his help in identifying specimens. Text and code was revised by ChatGPT-4. This work is supported in part by funds from the National Science Foundation (NSF: IRES#2246236). S.B. acknowledges funding support from NSF Grants CAREER iOS-1941933. The authors also thank the Regional Forestry Service in Madre de Dios, Peru (Collection Permit: 1288-2022-GOREMAD-GRFYFS.)

## Notes

### Competing Interest Statement

The authors have declared no competing interest.

### Summary of Updates

1. Revised Narrative - We have broadly softened the language around conclusions involving hexapods and instead focus on katydids (Tettigoniidae). We included additional references in the introduction and discussion to broaden the scope of the manuscript and provide relevant comparisons to other insect gait studies in the literature. 2. Additional Analyses - a. We conducted new mixed models where we included the mass of the individual katydid as a fixed effect for both the LMM on speed and the GLMM on gait. We included revised and updated tables (Tables 1 and 2) that show the fixed effects and their interactions for both models. b. We manually tracked a subset of the climbing videos (n=45 trials). From these tracks, we calculated the average stride length, average stance duration, and the average duty factor for katydids to quantify the stability of stance in relation to angle and mass. We included an additional panel (Figure 4e) and additional figures in the supplemental materials (S.Figs. 3-5). 3. New Results - a. Mass was found to have a large effect on both walking speed and gait preference. b. Substrate roughness has no significant effect on gait preference in the updated models. c. Duty factor increases with both the incline angle and insect mass. 4. Supplementary Additions - a. We included new measurements and a supplemental figure to calculate the observed power of our models, as requested by Reviewer 1. b. We include a new figure on duty factor and how it relates to the scored gait behavior, to show how consistent the authors were in gait classifications. c. We include additional supplemental tables on the LMM and GLMM, including: i. Comparisons between 2-way and 3-way models (S. Table 1,4) ii. Comparisons on whether random intercept and random slope is necessary in models. (S. Table 2) iii. Random effects for the LMM and GLMM (S. Table 3,5) d. Additional panels on the supplemental figures showing the residuals of the LMM and GLMM (S. Fig 6,7).

