## Supplemental Information for "Katydids Shift to Higher-Stability Gaits When Climbing Inclined Substrates"

### Supplemental Information for Riiska et al.,

**Supplemental Video 1.** A characteristic trial of a katydid individual using an alternating tripod gait to climb. The individual ID is 15, the incline angle is 60˚, and the sandpaper grit is 120.

**Supplemental Video 2.** A characteristic trial of a katydid individual using a metachronal gait to climb (i.e., not tripod). The individual ID is 15, the incline angle is 90˚, and the sandpaper grit is 40.

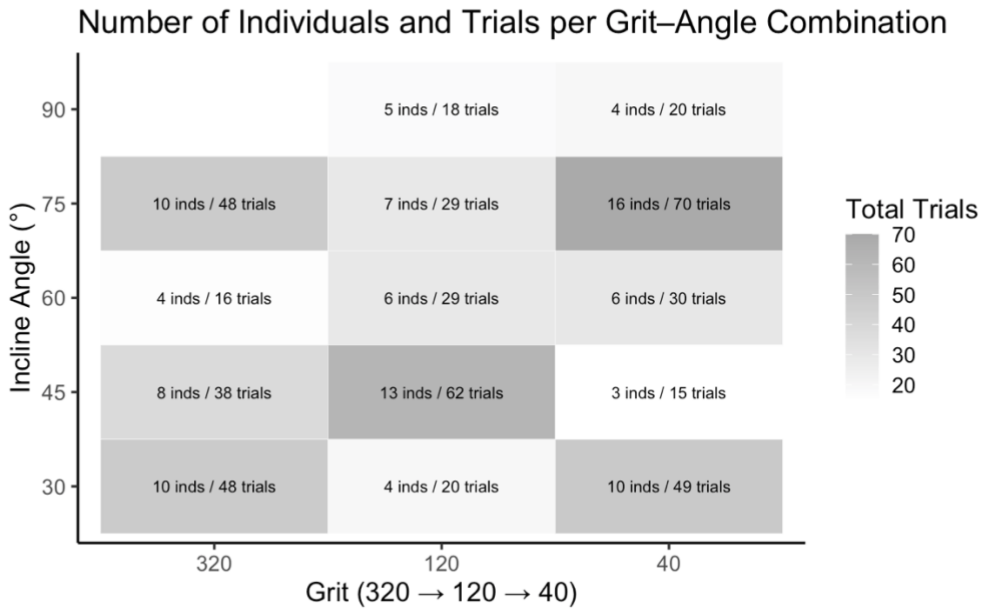


**Supplemental Figure 1**. The number of trials and number of individuals within each of the distinct combinations of grit and incline angle. There was a total of 492 distinct climbing trials across 24 individuals.

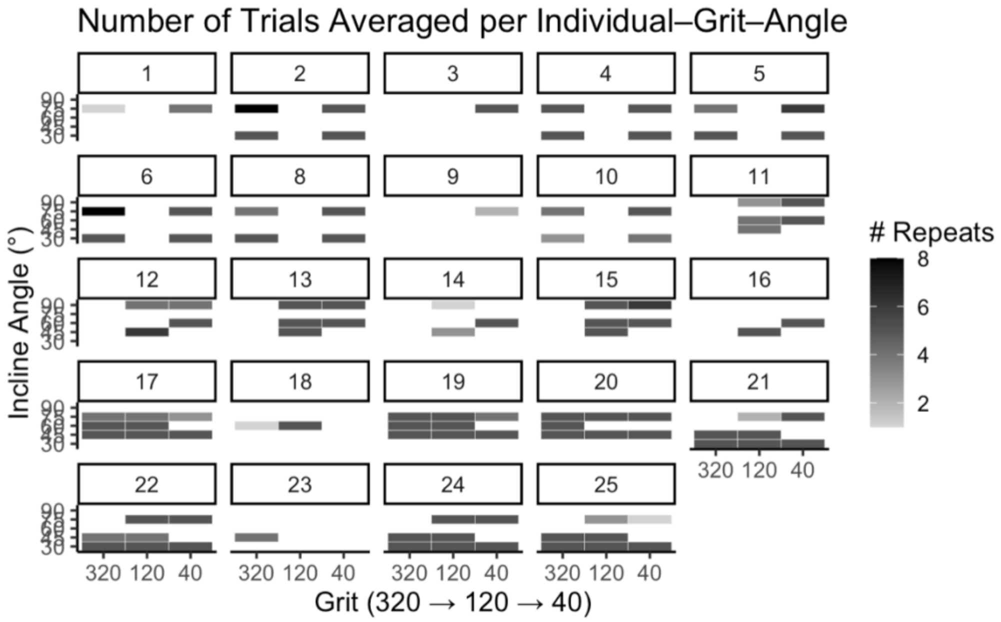


**Supplemental Figure 2**. The number of trials for each of the 24 individuals and the corresponding incline and grit.

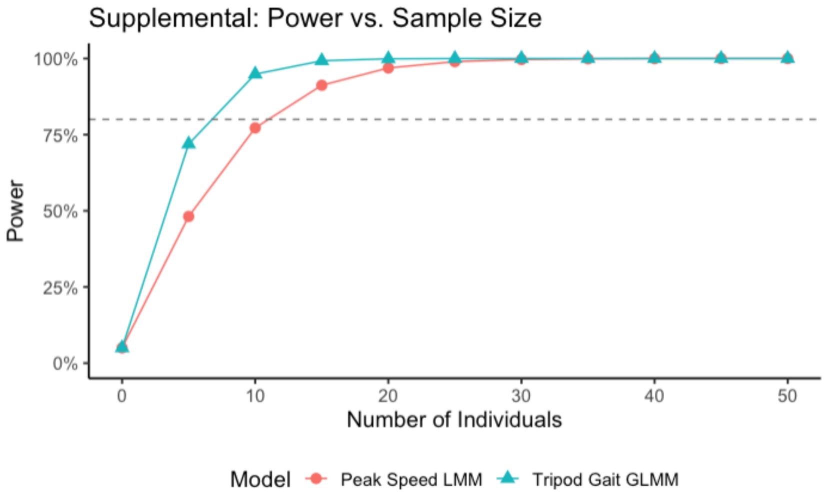


**Supplemental Figure 3**. The power as a function of sample size (n=0 - 50) for detecting the effect of incline angle on walking speed and tripod gait usage. Curves are computed from the observed effect sizes and standard errors from the final models run in this dataset with each point representing the power that a two-sided test would reject the null if the true effect equaled the observed effect.

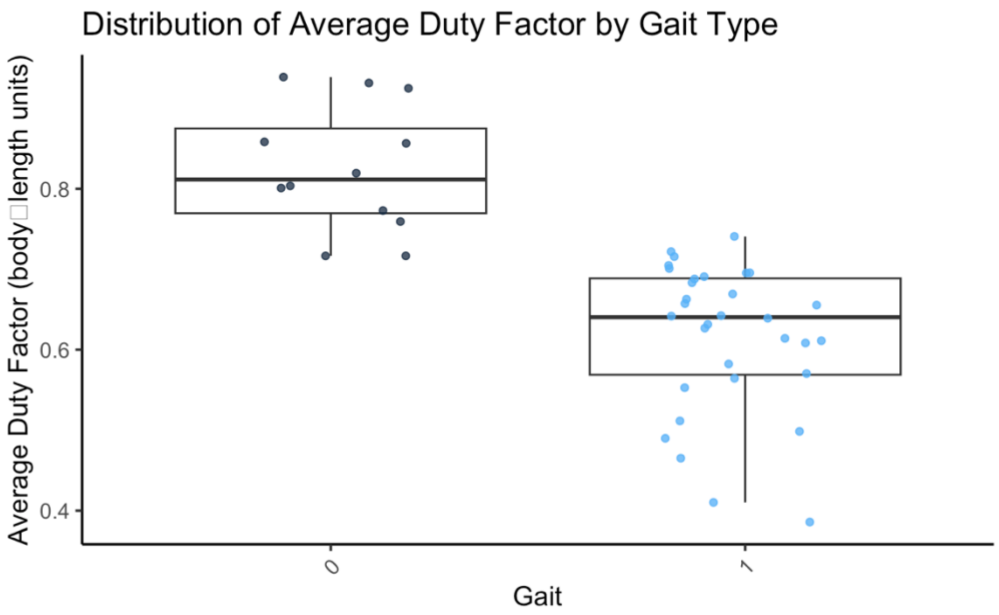


**Supplemental Figure 4.** The relationship between Duty Factor and the assigned gait shows that behavioral scoring was consistent in how gait related to quantified gait biomechanics. The 0 represents a non-tripod gait, and the 1 represents an alternating tripod gait.

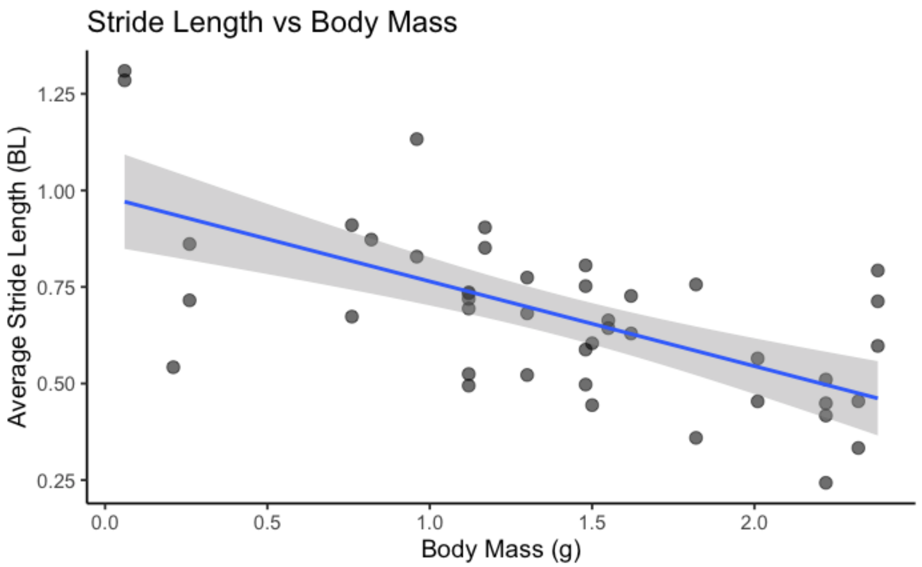


**Supplemental Figure 5. The a**verage stride length of katydids decreases with increasing body size. Points represent the average stride length across limbs of an individual katydid for a given substrate and grit combination. The blue line represents a linear regression, with grey shading indicating the standard deviation.

**Linear Mixed Model:**

Models:

LMM_2way: PeakSpeed_BLps ~ (grit_c + angle_c + Mass_c)^2 + (1 | Individual)

LMM_3way: PeakSpeed_BLps ~ grit_c * angle_c * Mass_c + (1 | Individual)

**Supplemental Table 1.** Comparisons between a 2-way and a 3-way linear mixed model shows that a three-way model (which includes a fixed effect has a statistically stronger fit. However, when using MuMIn::dredge() the top-ranked subset included two-way terms due to rank-deficiency warnings from three-way model. Because the two-way model yielded stable estimates and the interpretation was more straightforward we opted for the more parsimonious model.

| | npar | AIC | BIC | logLik | deviance | Chisq | Df | Pr(>Chisq) |
|---|---|---|---|---|---|---|---|---|
| LMM_2way: | 9 | 332.18 | 356.15 | -157.09 | 314.18 | | | |
| LMM_3way: | 10 | 327.51 | 354.14 | -153.75 | 307.51 | 6.6724 | 1 | 0.009792 ** |

**Supplemental Table 2.** Linear mixed model comparison for whether to include a random intercept or a random intercept with a random slope model shows that a random intercept is sufficient.

| | npar | AIC | BIC | logLik | deviance | Chisq | Df | Pr(>Chisq) |
|---|---|---|---|---|---|---|---|---|
| LMM_RI_only | 9 | 332.18 | 356.15 | -157.09 | 314.18 | | | |
| LMM_RS | 11 | 333.40 | 362.70 | -155.70 | 311.40 | 2.7822 | 2 | 0.2488 |

**Supplemental Table 3.** The random effects for the linear mixed model output for peak speed.

**Random-Effects (LMM of Peak Speed)**

| **Grouping Factor** | **Variance** | **Std. Dev.** |
|---|---|---|

| | | |
|---|---|---|
| Individual (Intercept) | 1.2871 | 1.1345 |
| Residual | 0.7714 | 0.8783 |

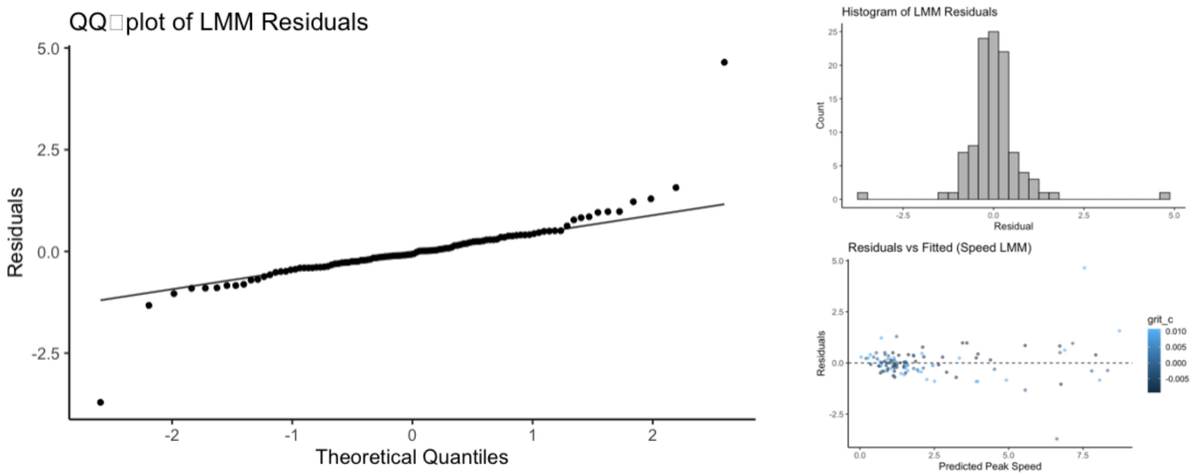


**Supplemental Figure 6.** (A) a Q-Q plot showing residuals of the average peak linear speed LMM model. (B) A histogram of rhe residuals show they predominantly fall near zero. (C) The residuals against the predicted peak speed.

**Generalized Linear Mixed Model:**

Models:

GLMM_2way: cbind(successes, failures) ~ (grit_c + angle_c + Mass_c)^2 + (1 | Individual)

GLMM_3way: cbind(successes, failures) ~ grit_c * angle_c * Mass_c + (1 | Individual)

**Supplemental Table 4.** Comparisons between a 2-way and a 3-way generalized linear mixed model (GLMM) shows that a two-way model (which only include the two-way interactions between fixed effects) provides a stronger fit to the data.

| | | npar | AIC | BIC | logLik | deviance | Chisq | Df | Pr(>Chisq) |
|---|---|---|---|---|---|---|---|---|---|
| glmm_gait_2way | 8 | 302.65 | 323.96 | -143.32 | 286.65 | | | | |
| glmm_gait_3way | 9 | 303.62 | 327.59 | -142.81 | 285.62 | 1.0312 | | 1 | 0.3099 |

**Supplemental Table 5.** The random effects for the generalized linear mixed model (GLMM) output for peak speed.

**Random-Effects (GLMM of Tripod Gait)**

| Grouping Factor | Term | Variance | Std. Dev. |
|---|---|---|---|
| rowID | (Intercept) | 5.2470 | 2.2906 |
| Individual | (Intercept) | 1.1565 | 1.0754 |
| Individual | Angle | 0.9029 | 0.9502 |

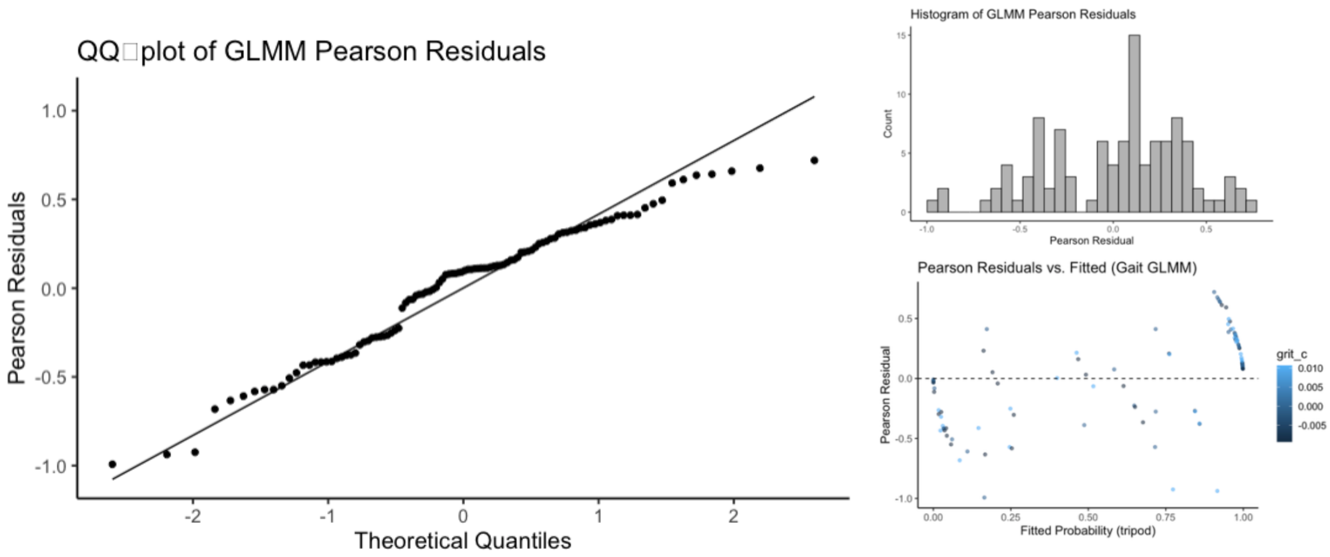


**Supplemental Figure 7.** (A) a Q-Q plot showing residuals for the gait probability GLMM. (B) A histogram of the residuals shows they have a relatively wide spread around 0. (C) The residuals against the predicted peak speed.

**Supplemental Note 1:** Species Identifications by Dr. David A. Nickle [retired]
Systematic Entomology Laboratory, USDA

1. adult, female: (Orthoptera: Tettigoniidae: Pseudophyllinae)
   *Teleutias* species possibly *fasciatus* Brunner von Wattenwyl

2. nymph, male: (Orthoptera: Tettigoniidae: Pseudophyllinae)
   *Schedocentrus* species

3. nymph, female: (Orthoptera: Tettigoniidae: Pseudophyllinae)
   possibly*Teleutias* species

4. nymph, female: (Orthoptera: Tettigoniidae: Pseudophyllinae)

5. nymph, female: (Orthoptera: Tettigoniidae: Pseudophyllinae)

6. adult, male: (Orthoptera: Tettigoniidae: Pseudophyllinae)

8. adult, female: (Orthoptera: Tettigoniidae: Phaneropterinae)

9. adult, female: (Orthoptera: Tettigoniidae: Phaneropterinae)

10. adult, female: (Orthoptera: Tettigoniidae: Pseudophyllinae)

11. adult, female: (Orthoptera: Tettigoniidae: Pseudophyllinae)
    *Cycloptera specularis* (Burmeister)

12. adult, female: (Orthoptera: Tettigoniidae: Pseudophyllinae)
    *Leptotettix voluptarius distinctus* Beier

13. adult, female: (Orthoptera: Tettigoniidae: Pseudophyllinae)

14. adult, female: (Orthoptera: Tettigoniidae: Pseudophyllinae)

15. nymph, male: (Orthoptera: Tettigoniidae: Pseudophyllinae)
    *Choeroparnops* species

16. nymph, male: (Orthoptera: Tettigoniidae: Pseudophyllinae)

17. adult, male: (Orthoptera: Tettigoniidae: Pseudophyllinae)

18. adult, male: (Orthoptera: Tettigoniidae: Pseudophyllinae)

19. adult, female: (Orthoptera: Tettigoniidae: Pseudophyllinae)

20. nymph, female: (Orthoptera: Tettigoniidae: Pseudophyllinae)

21. adult, female: (Orthoptera: Tettigoniidae: Pseudophyllinae)
*Leptotettix* species

22. adult, female: (Orthoptera: Tettigoniidae: Pseudophyllinae)

23. adult, male: (Orthoptera: Tettigoniidae: Conocephalinae)
*Copiphora gracilis* Scudder

24. nymph, male: (Orthoptera: Tettigoniidae: Pseudophyllinae)

25. (Orthoptera: Tettigoniidae: Phaneropterinae)
*Steirododon* species

Identifications by David A. Nickle [retired]
Systematic Entomology Laboratory, USDA
